# Functional connectivity differences in adult’s ADHD – a MEG study

**DOI:** 10.1101/2025.07.09.663963

**Authors:** Pedram Mouseli, Benjamin T. Dunkley, Massieh Moayedi

## Abstract

The neurobiology of adult Attention-Deficit/Hyperactivity Disorder (ADHD), particularly functional brain network connectivity, remain poorly understood. Magnetoencephalography (MEG) can reveal frequency-specific network dynamics given its high temporal resolution. Here, we investigated intrinsic functional connectivity differences between adults with ADHD (n = 24) and healthy controls (n = 44) using MEG data from the Open MEG Archive (OMEGA) dataset. We employed source reconstruction (Destrieux atlas), weighted phase-lag index (wPLI) connectivity across six frequency bands, graph theory metrics (Characteristic Path Length (CPL), node strength, clustering coefficient), and network-based statistics (NBS). We observed widespread hypo-connectivity in the high-gamma band (50-150 Hz) in adults with ADHD compared to controls. NBS identified a significant high-gamma sub-network, predominantly involving dorsal and ventral attention networks (DAN/VAN) and default mode network (DMN) nodes centered around a left fusiform gyrus hub, where all constituent connections exhibited consistently lower connectivity in the ADHD group. Globally, this was reflected in reduced gamma network integration (longer CPL) within the DAN and VAN. Locally, reduced high-gamma clustering was observed in VAN nodes (e.g., insula) and reduced node strength in a DAN region (postcentral sulcus). Predictive modeling using ElasticNet regression confirmed the importance of high-gamma metrics, with CPL measures yielding moderate classification accuracy (AUC ≈ 0.70–0.73). In contrast to high-gamma findings, the alpha band showed increased integration (shorter CPL) within the DMN and between the VAN and DAN in the ADHD group, alongside differences in alpha and beta band node properties in cingulate and somatomotor regions. Our findings reveal robust, frequency-dependent functional network alterations in adult ADHD, particularly highlighting disrupted high-frequency communication within and between key cognitive networks.

## 1. INTRODUCTION

Attention-Deficit/Hyperactivity Disorder (ADHD) is a prevalent neurodevelopmental disorder characterized by developmentally inappropriate levels of inattention, impulsivity, and hyperactivity. Affecting approximately 5% of school-aged children (Polanczyk et al., 2007) and persisting into adulthood in a significant number of cases (Mannuzza et al., 2003), ADHD poses a considerable burden on individuals, their families, and society (Barkley, 2002; Biederman et al., 1991; Matza et al., 2005). While much research has focused on ADHD in childhood, there is a growing recognition of the need to understand the condition in adults, where it is often underdiagnosed and can present with different symptom profiles (Asherson et al., 2012; Fayyad et al., 2017; Simon et al., 2009). Furthermore, even though ADHD is estimated to persist in as many as 65% of cases into adulthood (Mannuzza et al., 2003), the neurobiology of adult ADHD remains poorly understood.

The neurobiological basis of ADHD is thought to be related to widespread brain structure and function abnormalities, and may be better conceptualized as disrupted connectivity within and between large-scale brain networks (Konrad & Eickhoff, 2010). Crucially, the dynamic organization of these networks is thought to be mechanistically subserved by neural synchronization, where rhythmic, phase-aligned oscillations coordinate neural firing to selectively gate information flow and modulate synaptic efficacy (Fries, 2005, 2015). This “communication through coherence” allows for the precise temporal coordination necessary for efficient information integration and segregation—processes vital for cognitive functions typically impaired in ADHD. Investigating these rapid neural dynamics, particularly frequency-specific synchronization patterns, is therefore essential for a deeper mechanistic understanding of adult ADHD. Studies investigating spontaneous brain network activity in ADHD have reported several key networks in ADHD pathophysiology: the fronto-striatal network (Bush et al., 2005; Cubillo et al., 2012; Sun et al., 2012), which is crucial for executive functions, attention, and decision-making; the default mode network (DMN) (Buckner et al., 2008; Kucyi et al., 2015; Sonuga-Barke & Castellanos, 2007; Sun et al., 2012), which is active during rest and internally focused cognitive processes, and is prone to interfere with task-related cognitive processes when not appropriately suppressed during tasks; and the cognitive control (frontoparietal) network (Cai et al., 2021; Vincent et al., 2008; L. Wang et al., 2009; Zanto & Gazzaley, 2013; Zhang et al., 2024), which is essential for goal-directed behavior and cognitive flexibility. Furthermore, dorsal and ventral attention networks, which are critical for selective and sustained attention (Castellanos & Proal, 2012; Cortese et al., 2012; Fox et al., 2006; Y. Wang et al., 2025), have also been implicated. However, connectivity differences within and between these networks in ADHD compared to neurotypical controls have been mixed: studies report both hypo- and hyper-connectivity in ADHD (Sripada et al., 2014; van Rooij et al., 2015). Adding further complexity, evidence indicates that these connectivity differences can vary across the lifespan (Liu et al., 2023).

Despite these advances, considerable limitations and knowledge gaps persist concerning the precise neural mechanisms of network dysfunction in ADHD. While broad network abnormalities have been identified in adult ADHD, the specific nature of disruptions within these networks, from the level of individual nodes and connections to whole-brain and resting-state network properties, remains poorly understood. Crucially, the role of neural oscillatory activity, such as the high gamma band, in these network disruptions is under-explored, largely due to methodological constraints. Magnetoencephalography (MEG) offers superior temporal resolution, enabling the investigation of fast neural dynamics, including high-frequency oscillations like gamma activity, which are considered vital for cognitive processing.

The majority of existing studies have focused on investigating univariate edge-level differences in ADHD, with a less frequent focus on alterations at the local and network levels in terms of node-specific and network-specific properties. Graph theory provides a powerful framework to characterize these more granular network properties. Global network topology can be assessed using metrics like the characteristic path length (CPL), which quantifies the average shortest communication pathway between all nodes, reflecting the overall efficiency of information integration across the brain. As such, a longer CPL suggests a less integrated network, while a shorter CPL may indicate an overly integrated or less segregated network configuration, with either state potentially impacting cognitive function. At a more local level, nodal properties provide insight into the roles of specific brain regions. The clustering coefficient measures the degree of local interconnectivity of a node, indicating the efficiency of local information processing, while node strength reflects a region’s overall functional influence. Alterations in these metrics within specific resting-state networks (RSNs) could pinpoint key dysfunctional hubs contributing to ADHD pathophysiology. Furthermore, examining the organization of interconnected edges, rather than isolated univariate differences, can reveal specific sub-networks of disrupted connectivity.

Furthermore, while much of the existing neuroimaging research in ADHD has focused on pediatric populations, a dedicated investigation into adult ADHD is warranted, given that the disorder can present with distinct neurobiological features and symptom profiles in adulthood. Addressing these gaps is crucial, as a more detailed understanding of network dysfunction can inform targeted therapeutic interventions, enhance comprehension of the neurobiological underpinnings of its atypical presentations and high comorbidity rates, and help elucidate the mechanisms underlying ADHD heterogeneity.

The current work addresses these needs by adopting a granular approach to investigate functional connectivity in adult ADHD, examining specific node properties, edge-level differences, and network-level interactions across different frequency bands. Specifically, our research aims to: (1) determine whether there are group differences between adults with ADHD and neurotypical adults in: (1) CPL within and between RSNs, (2) RSN nodal network properties ; and (3) whole-brain connectivity networks at the edge-level across various frequency bands. Specifically, this research focuses on the role of different frequency bands in network dysfunction to provide a more nuanced characterization of the neurophysiological differences associated with adult ADHD.

## 2. MATERIALS AND METHODS

This study investigated functional connectivity differences in adults with ADHD using MEG data from the publicly available OMEGA dataset (Niso et al., 2016). The dataset comprises resting-state MEG recordings with ECG and EOG signals recorded simultaneously, alongside structural MRI, from 68 participants: 24 adults diagnosed with ADHD (15 female, mean age 21.25 ± 4.44 years) and 44 healthy controls (28 female, mean age 22.29 ± 2.01 years). All participants provided informed consent as part of the original OMEGA data collection protocol. This study was also reviewed and approved by the University of Toronto Human Research Ethics Board (protocol # 00045243).

### 2.1 Data

Resting-state MEG data were acquired using a 275-channel axial gradiometer system (CTF MEG, Coquitlam, BC). Participants were instructed to remain still with eyes open for a 4 to 5-minute duration. As most resting-state recordings obtained from the ADHD group were 4 minutes, MEG data were truncated to the initial 4 minutes to ensure uniformity in data length across all participants. Simultaneous electrophysiological recordings, including electrocardiogram (ECG) and vertical and horizontal electrooculogram (EOG), were obtained to monitor cardiac activity and eye movements, respectively. Structural T1-weighted magnetic resonance images (MRI) were acquired for each participant and used to facilitate source localization. MEG data were recorded at a sampling rate of 2,400 Hz and low-pass filtered at 600 Hz during acquisition. Multiple resting-state runs were available for most participants, but for the primary analyses, only data from the first run were utilized to maintain consistency across participants. However, for the training of predictive models, all available runs (106 control, 126 ADHD runs total) were incorporated to maximize training sample size and model robustness.

### 2.2 MEG Data Preprocessing

Data underwent a comprehensive preprocessing pipeline using established best practices in MEG research (Gross et al., 2013) and implemented in the Brainstorm toolbox (Tadel et al., 2011). Initial preprocessing steps aimed to mitigate noise and artifact contamination. Line noise at 60 Hz and its harmonics (120 Hz and 180 Hz) were attenuated using a notch filter. A high-pass filter with a cutoff of 0.3 Hz was applied to remove slow-drift and low-frequency artifacts. To address physiological artifacts, heartbeat, blink, and saccade events were identified using ECG and EOG signals, initially through automated detection algorithms and subsequently verified and refined through manual review. Signal-Space Projection (SSP) was then employed to remove signal components associated with these identified artifactual events, effectively reducing noise contributions from cardiac and ocular sources (Tadel et al., 2019; Tesche et al., 1995; Uusitalo & Ilmoniemi, 1997). Muscle artifact, specifically noise from neck muscle tension, was targeted by analyzing MEG signals in the 40-150 Hz range. Components reflecting muscle activity, showing concentrated power in posterior sensors close to neck muscles, were identified and removed. Following these automated and semi-automated preprocessing steps, MEG data were visually inspected for residual artifacts. Segments of data exhibiting poor signal quality, as well as malfunctioning channels identified through power spectrum density analysis, were manually excluded from further analysis.

### 2.3 Source Reconstruction and Functional Connectivity Analysis

Source reconstruction was performed to estimate neuronal activity at the cortical level. T1-weighted MRI data were processed using FreeSurfer v6.0.0 (Fischl, 2012) to segment and parcellate the cortex into regions based on the Destrieux atlas (Destrieux et al., 2010), which divides each hemisphere into 74 regions, resulting in a total of 148 cortical regions or nodes. FreeSurfer was also used to generate subject-specific head models for source estimation. Coregistration of MRI data with MEG sensor locations was achieved using digitized head points collected during each MEG recording session. Forward models were computed using the overlapping spheres method. Cortical source models were then constructed using Linearly Constrained Minimum Variance (LCMV) beamforming, implemented within the Brainstorm toolbox (Tadel et al., 2011). To mitigate the influence of varying source depths, covariance regularization was applied with a regularization parameter of 0.1. MEG source orientations were constrained to be normal to the cortical surface, with source activity estimated at approximately 15,000 dipoles distributed across the cortex.

Functional connectivity between cortical regions was then computed. For each participant, the average time series of dipole activity within each of the 148 Destrieux atlas regions was calculated, as is the default in Brainstorm v190508 (Brkić et al., 2023). For frequency domain analysis, data were segmented into four-second epochs, and frequency components were calculated using a multitaper fast Fourier transform (FFT) with spectral smoothing. Pairwise functional connectivity between these regional time series was estimated using the weighted phase lag index (wPLI) (Vinck et al., 2011). The wPLI was chosen as a measure of functional connectivity because it is relatively insensitive to the effects of volume conduction and signal leakage, minimizing spurious connectivity estimates. Whole-brain functional connectivity matrices were constructed across six predefined frequency bands: delta (1-4 Hz), theta (4-8 Hz), alpha (8-13 Hz), beta (13-30 Hz), gamma (30-50 Hz), and high gamma (50-150 Hz), representing pairwise wPLI values between all 148 regions. The absolute value of wPLI was used for subsequent graph theoretical analyses. The FieldTrip toolbox (Oostenveld et al., 2011) was used to perform all functional connectivity analyses.

### 2.4 Graph Theory Analysis

Graph theory analysis was employed to characterize the topological organization of the functional brain networks derived from the wPLI connectivity matrices. Brain regions were considered as nodes, and their functional connections (wPLI values) as edges in the graph. Both global and local graph measures were computed using the Brain Connectivity Toolbox (Rubinov et al., 2009), implemented in Python.

#### 2.4.1 Global Graph Analysis: Characteristic Path Length

Global network integration was assessed using characteristic path length (CPL). CPL represents the average shortest path length between all pairs of nodes in the network. CPL was calculated for the whole-brain network and also separately for within each of seven RSNs (Visual, Somatomotor, Dorsal Attention, Ventral Attention, Limbic, Frontoparietal, Default) as defined by Thomas Yeo et al. (2011). Regions from the Destrieux atlas were assigned to these seven RSNs based on majority voting, adapted from Sareen et al. (2021). Shortest paths were computed in the weighted network by inverting the connectivity values, such that higher connectivity corresponded to a shorter path, using Dijkstra’s algorithm (Dijkstra, 1959). Additionally, the average shortest path between RSNs was calculated by averaging the shortest paths between all nodes in one network to all nodes in another network (See Figure S1).

#### 2.4.2 Local Graph Analysis: Node Properties

Local graph measures were calculated to characterize the properties of individual brain regions within the network. These included: node strength (sum of weights of connections to a node), betweenness centrality (extent to which a node lies on shortest paths between other nodes), and clustering coefficient (tendency of a node’s neighbors to be interconnected).

### 2.5 Statistical Analysis

Group differences in global and local graph measures between ADHD and control groups were statistically assessed using non-parametric Wilcoxon rank-sum tests. For local graph measures, false discovery rate (FDR) correction was applied to account for multiple comparisons across all the nodes in the network.

Network-based statistic (NBS) (Zalesky et al., 2010) were used to identify sub-networks exhibiting significant connectivity differences between the groups at the edge level in each frequency band. The significance of each connection (edge) was evaluated using the Wilcoxon rank-sum test. Connections with a p-value below a threshold (0.001 for whole-brain network; 0.05 for between-network distances) and sharing common nodes were clustered into sub-networks. The size of these sub-networks, defined by the number of edges, served as the test statistic. The statistical significance of identified sub-networks was determined using a permutation test with 5000 iterations, assessing the null hypothesis of no group difference in network connectivity.

### 2.6 Predictive Modeling

To evaluate the predictive utility of network measures, separate ElasticNet logistic regression models were developed for within-network characteristic path length and between-network average shortest paths in each of the six frequency bands. ElasticNet, which incorporates both L1 and L2 regularization, was chosen to balance model complexity and prevent overfitting. A nested cross-validation approach was implemented. Hyperparameter tuning (regularization coefficient and L1 ratio) was performed using a grid search within an inner 10-fold cross-validation loop. Model performance was then evaluated using an outer 5-fold cross-validation. Given the slight class imbalance in the dataset (106 control runs, 126 ADHD runs for predictive modeling), model parameters were optimized to maximize the F1-score, balancing precision and recall. Class weights were adjusted inversely proportional to class frequencies to further mitigate the impact of class imbalance. To prevent data leakage and ensure robust generalization, stratified group k-fold cross-validation was employed in both inner and outer loops, ensuring that data from the same participant were not split across training and testing sets, while maintaining class distribution ratios within each fold. Model performance was quantified using the area under the receiver operating characteristic (ROC) curve (AUC). Furthermore, to explore the feature space of between-network distances, a Partial Least Squares Discriminant Analysis (PLS-DA) model with four components was trained on the between-network average shortest path data in the high-gamma band. Feature loadings from the PLS-DA model were examined to gain insights into the contribution of different between-network paths to group classification.

## 3. RESULTS

### 3.1 Network-Based Statistics

To further dissect network differences, we applied network-based statistics (NBS) to the whole-brain connectivity matrices in each frequency band. A significant sub-network differentiating the ADHD and control groups was identified specifically in the high-gamma band (p=0.009) (Figure 5A). This sub-network comprised 24 connections, predominantly located within the left hemisphere (14 intra-hemispheric left, 2 intra-hemispheric right, and 8 inter-hemispheric). Regions within the dorsal attention, ventral attention, and default mode networks were prominently represented in this sub-network. The left fusiform gyrus emerged as a hub within this sub-network, displaying the highest number of connections, particularly with nodes belonging to the dorsal attention, ventral attention, and default mode networks. Across all identified connections in this sub-network, connectivity was consistently reduced in the ADHD group compared to the control group (Figure 5A). The most pronounced group differences were observed in two connections originating from the inferior temporal gyrus: one linking to the anterior segment of the circular sulcus of the insula and the other to the lateral occipito-temporal sulcus (Figure 5B). These connections exhibited the most substantial disruptions in the ADHD group within the high-gamma frequency band.

### 3.2 Global Graph Analysis

We first investigated the characteristic path length (CPL), a measure of network integration, across the whole brain and within seven established RSNs. Analysis of within-network CPL revealed frequency-specific alterations. In the alpha frequency band, the default mode network (DMN) exhibited a significantly increased CPL in the control group compared to the ADHD group (p=0.038) (Figure 2A). Conversely, in the high-gamma band, both the dorsal attention network (DAN) (p=0.041) and ventral attention network (VAN) (p=0.029) as well as the visual network (p=0.038) demonstrated significantly elevated CPL in the ADHD group relative to controls (Figure 2A). To evaluate the discriminatory capacity of within-network CPL, we trained ElasticNet models for each frequency band to classify individuals as either control or ADHD based on these metrics. The high-gamma frequency band yielded the most robust predictive performance, achieving an Area Under the Curve (AUC) of 0.70 (Figure 2C), indicating a moderate ability to differentiate between the groups based on within-network CPL in this frequency range.

**Figure 1:**
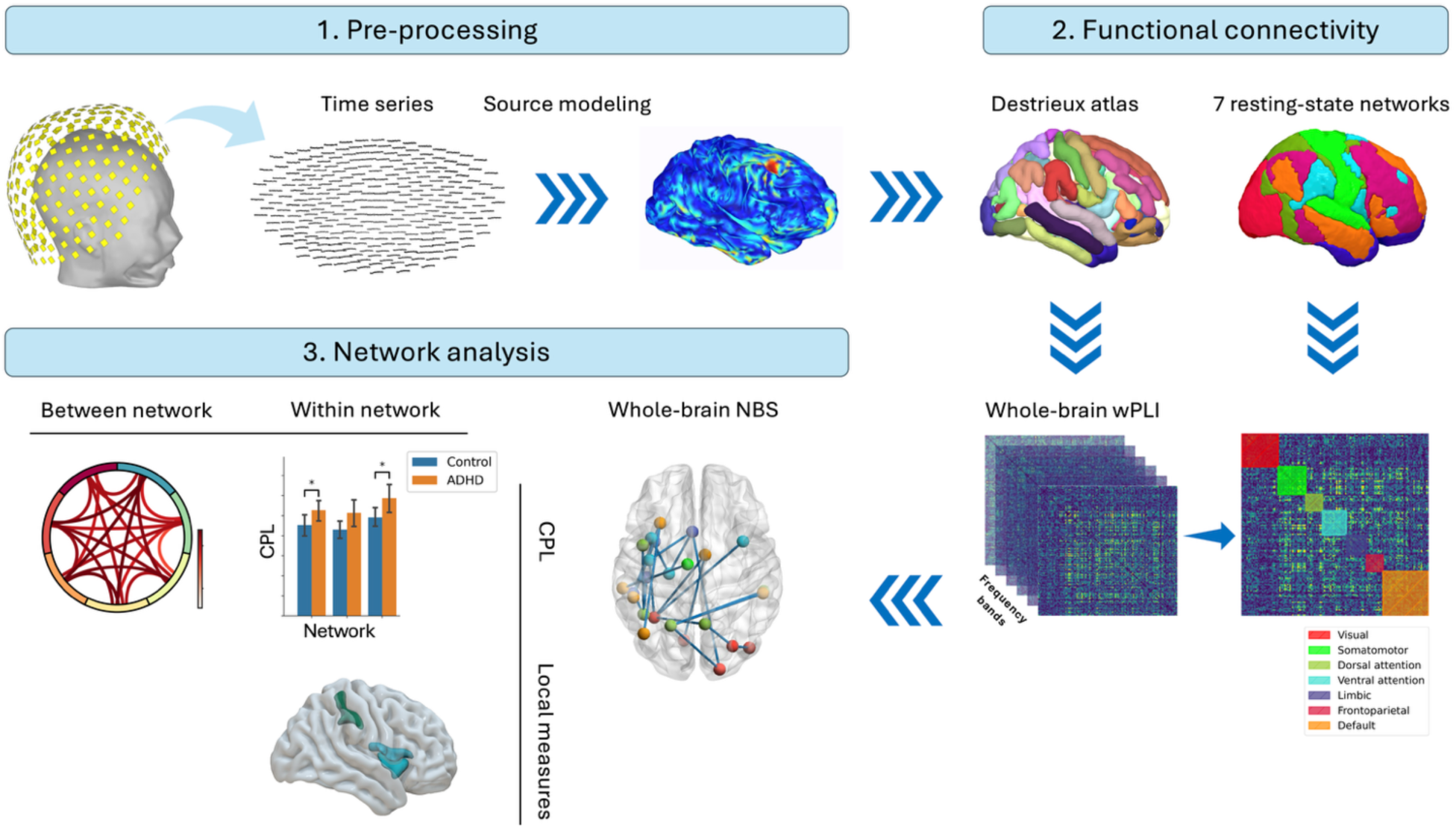
Methodological pipeline. MEG data underwent pre-processing and source modeling to extract cortical time series from 148 regions in the Destrieux atlas. Functional connectivity was calculated using wPLI across six frequency bands. Regions were assigned to 7 RSNs. Network analysis included calculating characteristic path length (CPL) within/between RSNs, local node measures, and network-based statistics (NBS) to identify significant whole-brain and between-network connectivity differences.

**Figure 2:**
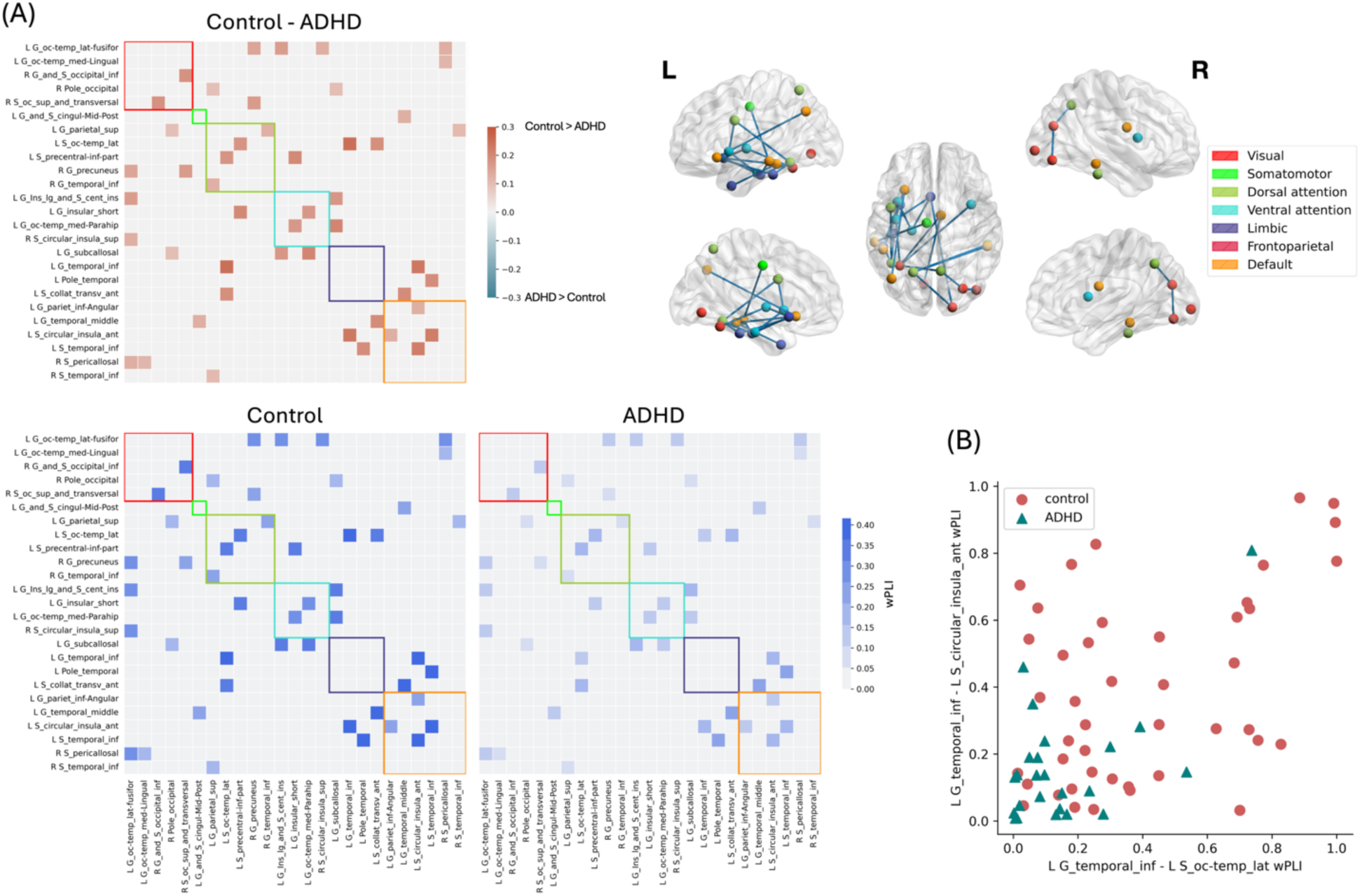
ADHD exhibit reduced high frequency synchrony compared to controls. (A) Sub-network identified using NBS in the high gamma band. This network consisted of 24 connections, with the majority located in the left hemisphere. Connectivity differences between the ADHD and control groups in the sub-network identified in the high gamma band showed with blue color bar. All connections in the ADHD group exhibited lower connectivity compared to the control group. The left fusiform gyrus had the highest number of connections, linking to regions in the dorsal attention, ventral attention, and default mode networks. (B) Strongest connections in the sub-network. Two connections, from the inferior temporal gyrus to the anterior segment of the circular sulcus of the insula and to the lateral occipito-temporal sulcus, showed the largest differences between the ADHD and control groups. These connections were notably disrupted in the ADHD group in the high gamma band.

Further global analysis extended to between-network connectivity, assessing the average shortest path between all pairs of the seven RSNs. Network-based statistics (NBS) identified a significant sub-network of between-network average shortest paths in the alpha band (p=0.041) (Figure 3A). This sub-network, characterized by a dorsal attention network hub, showed a consistent pattern of longer average shortest paths in the control group compared to the ADHD group. The most pronounced difference within this sub-network was observed in the average shortest path between the dorsal and ventral attention networks. To examine the predictive utility of between-network average shortest paths, we again employed ElasticNet models for each frequency band. Similar to the within-network CPL analysis, the high-gamma frequency band showed the highest classification accuracy based on all between-network CPL values, reaching an AUC of 0.73 (Figure 3C). A Partial Least Squares Discriminant Analysis (PLS-DA) classifier, trained with four components on the same high-gamma band data, was used to explore feature loadings. This analysis revealed that although most RSNs contributed strongly to the first component, the frontoparietal network accounted for much of the second component’s variance, the visual network for the third, and the ventral and dorsal attention networks for the fourth (Figure 3B).

**Figure 3:**
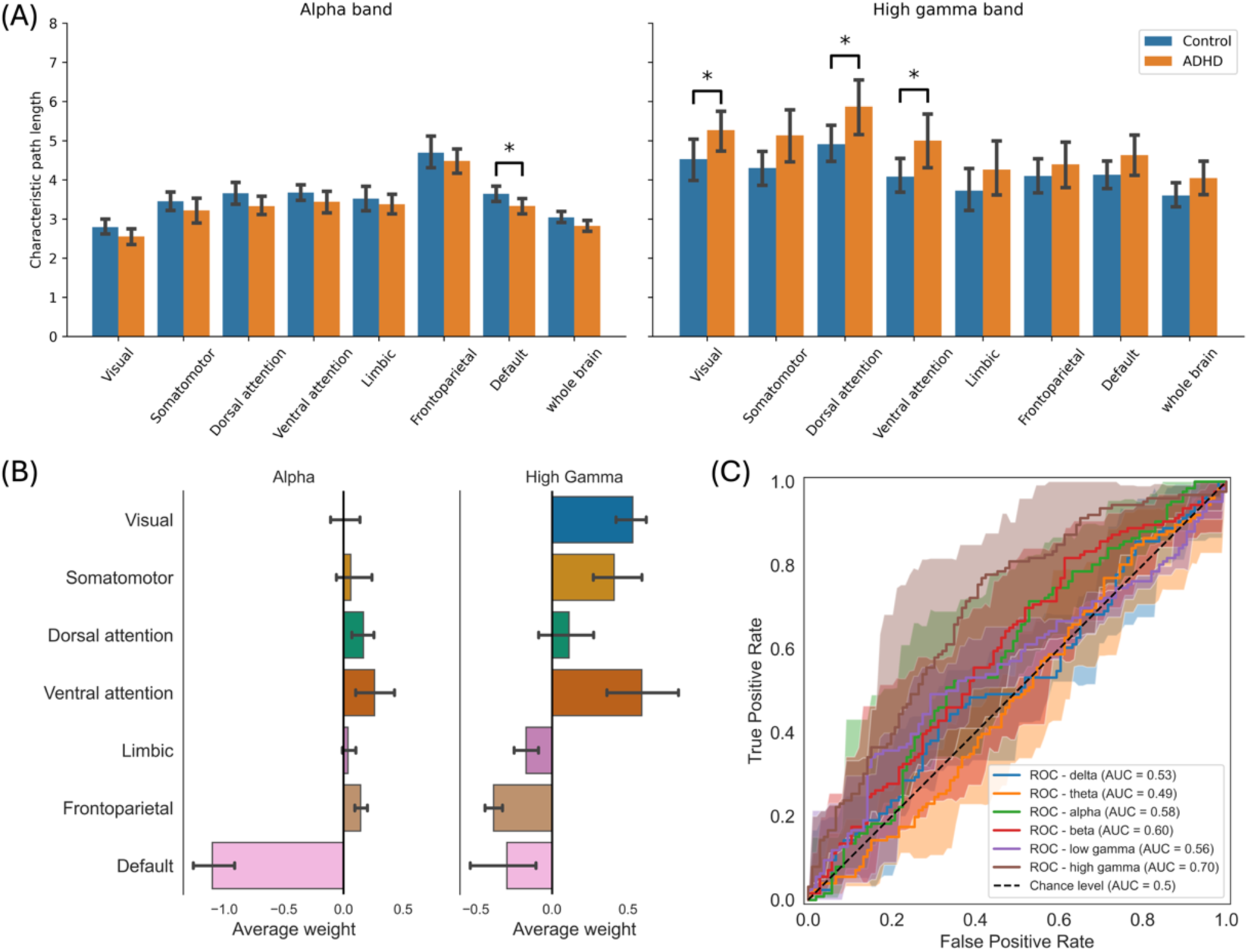
Global network integration differs in adult ADHD, with significant CPL changes observed in the alpha band for the Default Mode Network and in the high-gamma band for attention networks. (A) Characteristic path length of the functional connectivity network in the alpha and high gamma frequency bands. Significant differences between the ADHD and control groups are shown. No significant differences were observed in the other frequency bands. (B) Feature weights of the predictive models based on within network CPL in alpha and high-gamma frequency bands. (C) The receiver-operator curve (ROC) of ElasticNet models based on within network CPL in each frequency band. The shaded area represents the mean ± standard deviation of ROC curve from five folds.

### 3.3 Local Graph Analysis

Beyond global network properties, we investigated local graph measures to characterize the roles of individual brain regions within the functional networks. Specifically, we analyzed node strength, betweenness centrality, and clustering coefficient across frequency bands. Significant group differences in node strength and clustering coefficient were observed in the alpha, beta, and high-gamma frequency bands. In the alpha band, the middle-anterior part of the left cingulate gyrus and sulcus, a region within the ventral attention network, showed significantly increased node strength (p=0.008) and clustering coefficient (0.015) in the ADHD group compared to controls (Figure 4A). In the beta band, the right subcentral gyrus (central operculum) and sulci (p=0.025), along with the transverse temporal sulcus (0.025) within the somatomotor network, exhibited significantly greater node strength in the ADHD group (Figure 4B). Conversely, in the high-gamma band, the right postcentral sulcus, located within the dorsal attention network, demonstrated significantly reduced node strength (0.045) in the ADHD group. Furthermore, several regions within the ventral attention network showed significantly lower clustering coefficients in the ADHD group compared to controls, including the bilateral superior segment of the circular sulcus of the insula (p=0.036) and short insular gyri (p=0.036), as well as the left opercular part of the inferior frontal gyrus (IFG) (p=0.036) and the parahippocampal gyrus (p=0.036) (Figure 4C).

**Figure 4:**
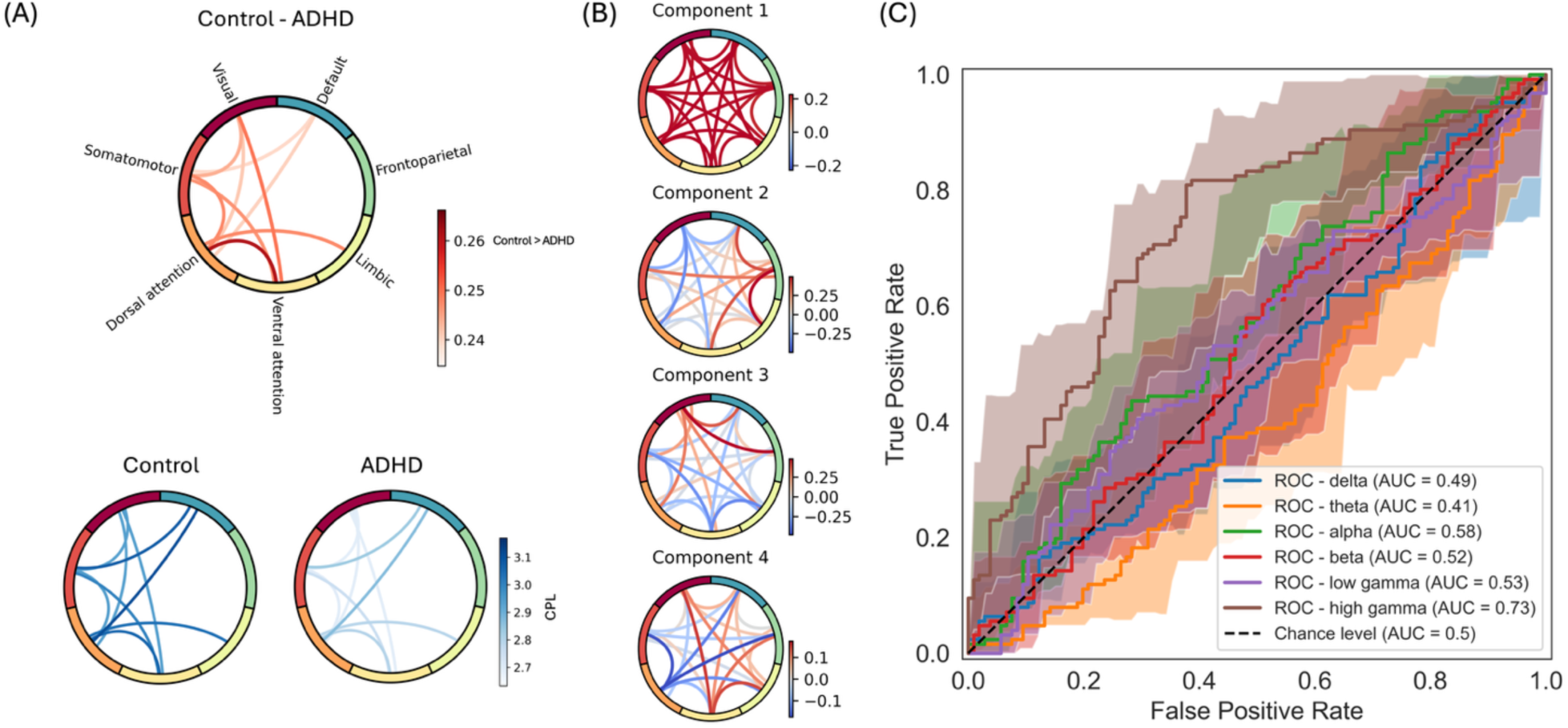
**A specific alpha-band sub-network, identified via NBS, is characterized by consistently longer average shortest paths across all its connections in controls relative to adults with ADHD**, **whereas high-gamma band paths demonstrate the most robust predictive power for differentiating the groups.** (A) An alpha-band sub-network derived from NBS on the average shortest paths between RSNs, showing consistently longer average shortest paths for all connections in the control group compared to the ADHD group. No significant sub-network was found in any other frequency band using NBS. (B) Feature loadings of a four component PLS-DA model classifying ADHD and control groups based on between network average shortest paths in high-gamma frequency band. (C) The ROC curve of ElasticNet models based on between network average shortest paths in each frequency band. The shaded area represents the mean ± standard deviation of ROC curve from five folds.

**Figure 5:**
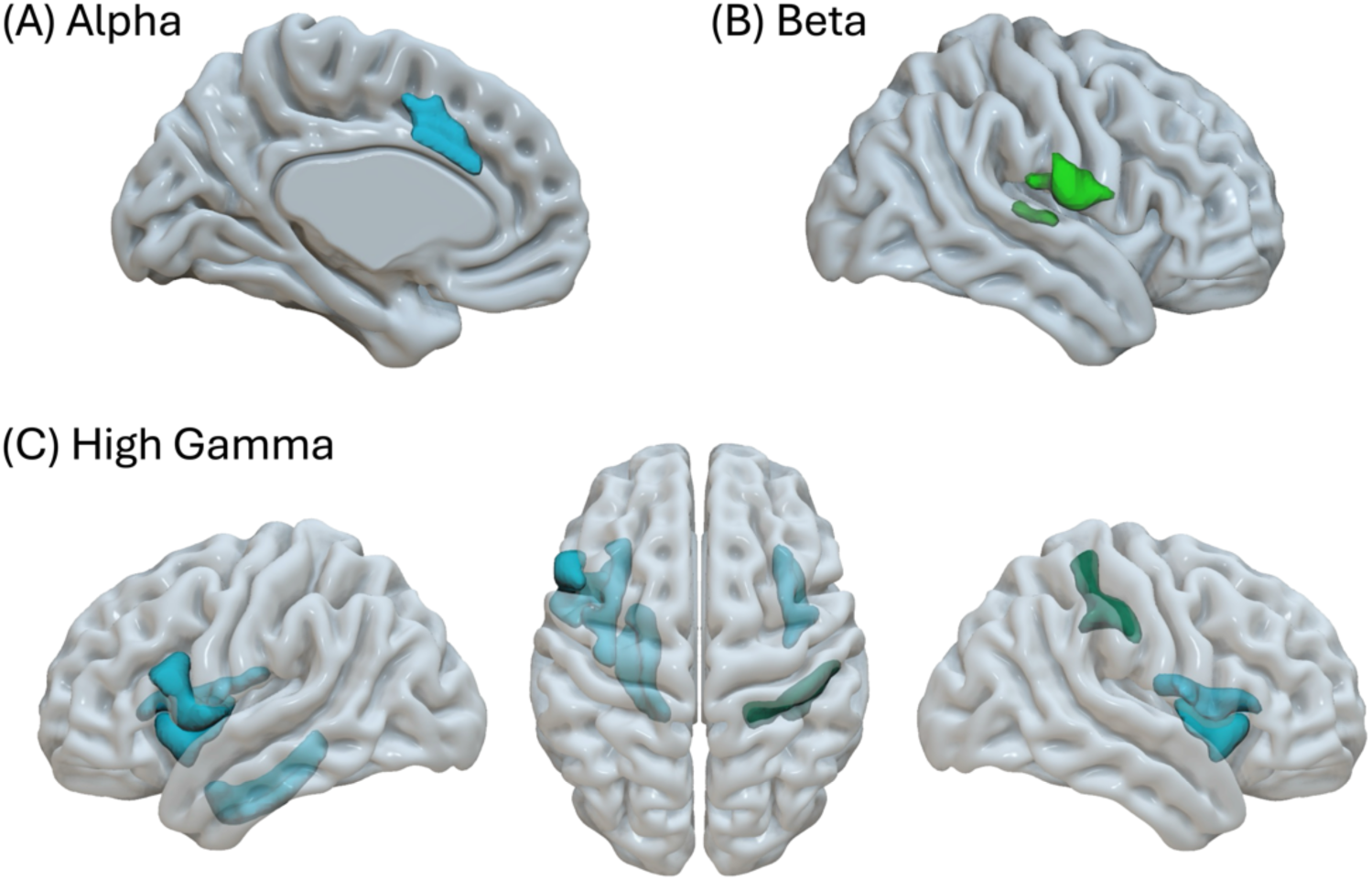
Specific brain regions within the ventral attention, dorsal attention, and somatomotor networks exhibit altered local connectivity properties (node strength, clustering) in adults with ADHD across different frequency bands. (A) Differences in local graph measures in the alpha band. The middle-anterior part of the left cingulate gyrus and sulcus in the ventral attention network showed significantly greater clustering coefficient and node strength in the ADHD group compared to controls. (B) Differences in local graph measures in the beta band. The right subcentral gyrus (central operculum) and sulci, along with the transverse temporal sulcus in the somatomotor network, had significantly greater node strength in the ADHD group compared to controls. (C) Differences in local graph measures in the high gamma band. The right postcentral sulcus within the dorsal attention network showed significantly lower node strength in the ADHD group. Several regions within the ventral attention network, including bilateral superior segment of the circular sulcus of the insula and short insular gyri, along with left opercular part of the IFG and parahippocampal gyrus exhibited significantly lower clustering coefficients in the ADHD group compared to controls.

## 4. DISCUSSION

### 4.1 Dominant High-Gamma Hypo-connectivity: A Core Neurophysiological Finding in Adult ADHD

Leveraging MEG’s high temporal resolution, this study investigated functional connectivity alterations in adults with ADHD using a granular, multi-frequency, multi-level network analysis. We found frequency-specific disruptions: key results include reduced high-gamma integration within dorsal/ventral attention networks and widespread hypo-connectivity in a high-gamma sub-network (centered on the left fusiform gyrus) involving attention/default mode networks. Conversely, increased alpha-band integration occurred within the DMN. Local analyses also revealed altered alpha, beta, and high-gamma node properties in DAN, VAN, and somatomotor regions.

To pinpoint specific connection differences, we applied network-based statistics (NBS) to edge-level weighted connectivity matrices. A significant sub-network differentiating the groups emerged exclusively in the high-gamma band. This sub-network consisted of 24 connections where connectivity was consistently lower in the ADHD group compared to controls. These hypo-connected edges were predominantly located within the left hemisphere and primarily involved nodes within the DAN, VAN, and DMN. The left fusiform gyrus emerged as a significant hub within this hypo-connected network, showing reduced connectivity particularly to nodes in the DAN, VAN, and DMN. The left fusiform gyrus emerged as a significant hub within this hypo-connected network, displaying reduced connectivity particularly to nodes in the DAN, VAN, and DMN. The fusiform gyrus is a key area for higher-level visual processing, particularly for objects and faces, and potentially visual mental imagery (Zhou et al., 2024). Notably, activity within the fusiform gyrus, including induced gamma-band oscillations (often peaking above 60 Hz), is strongly modulated by covert visual attention (Tallon-Baudry et al., 2005; Wojciulik et al., 1998). Increased attention typically enhances fusiform responses (Wojciulik et al., 1998) and stimulus-related gamma oscillations (Tallon-Baudry et al., 2005). Therefore, the widespread high-gamma hypo-connectivity observed here, linking the fusiform gyrus hub to major attention (DAN, VAN) and internal state (DMN) networks, may represent a significant deficit in integrating or modulating visual processing in adult ADHD. This reduced high-frequency synchrony could reflect impaired attentional gating of visual information processed within the fusiform, or difficulties coordinating its activity with the demands set by attentional control networks or the internal context represented by the DMN. The most substantial connectivity reductions were observed between the left inferior temporal gyrus and both the anterior segment of the circular sulcus of the insula and the lateral occipito-temporal sulcus. These pathways are potentially involved in integrating sensory information with salience/emotional processing (insula) and higher-level visual/semantic processing (occipito-temporal sulcus), suggesting these specific integrative functions may be particularly disrupted at high frequencies in adult ADHD.

Global network integration, assessed via characteristic path length (CPL), revealed that the ADHD group exhibited significantly longer CPL (reduced integration) in both DAN and VAN within the high-gamma band. This suggests less efficient information transfer within these core attention systems, potentially reflecting difficulties in processes reliant on high-frequency oscillations, such as maintaining attentional focus. This finding aligns with evidence linking gamma activity to selective attention and attentional network synchronization (Gruber et al., 1999; Ray et al., 2008; Doesburg et al., 2008), and with reports of reduced intrinsic gamma power in adult ADHD (Tombor et al., 2021). Such deficits could impair information transfer critical for attentional control.

Critically, the high-gamma band showed evidence of local processing deficits: the right postcentral sulcus (DAN) exhibited reduced node strength, while several VAN regions (bilateral superior segment of the circular sulcus of insula, short insular gyri, left opercular part of IFG, parahippocampal gyrus) showed significantly lower clustering coefficients in the ADHD group relative to controls. The insula, particularly the anterior portion, is a critical hub within the salience network (SN), implicated in detecting behaviorally relevant stimuli and initiating dynamic switching between large-scale networks like the default mode network and central executive network (Menon & Uddin, 2010; Nelson et al., 2010). Both structural (reduced volume) (Lopez-Larson et al., 2012) and functional connectivity alterations of the anterior insula have been previously reported in ADHD (Zhao et al., 2017). The clustering coefficient reflects the density of connections among a node’s neighbors, indexing the efficiency of local information processing. Our finding of reduced high-gamma clustering coefficient in insular regions (and associated opercular IFG) suggests less integrated or less efficient local processing within these key nodes of the salience/ventral attention system in adult ADHD, specifically in the high-frequency domain essential for fast neural communication. This local processing inefficiency could impair the insula’s ability to effectively perform its functions in saliency detection and network switching (Menon & Uddin, 2010), contributing to the attentional and regulatory deficits characteristic of ADHD. Similarly, the reduced high-gamma clustering in the parahippocampal gyrus, a region critical for memory processes but also interacting with attentional networks, points towards broader disruptions in local high-frequency network organization in adult ADHD, though its specific contribution requires further investigation. These findings collectively suggest diminished functional influence (strength) and less efficient local information processing (clustering) within key nodes of the attention networks at high frequencies, potentially contributing to the global integration deficits observed in the same frequency band.

The observed disruptions in high-gamma band connectivity gain significance when considering their putative underlying neural mechanisms. Gamma oscillations are widely thought to emerge from the interaction between excitatory principal cells and inhibitory interneurons (Buzsáki & Wang, 2012). In this ‘E-I’ (excitatory-inhibitory) loop model, pyramidal cells excite interneurons, which in turn provide rapid, precisely-timed feedback inhibition onto the pyramidal cells, creating a rhythmic cycle of activity (Bartos et al., 2007; Buzsáki & Wang, 2012). While this mechanism is well-established for lower gamma frequencies, higher gamma activity (>60 Hz), such as that observed in our study, may involve additional or distinct processes, including contributions from gap junctions and different characteristics of the underlying E-I balance (Uhlhaas et al., 2011). Therefore, the widespread high-gamma hypo-connectivity found in our ADHD group—spanning global integration, local processing, and specific edge-level connections—likely reflects a fundamental deficit in the efficacy or temporal precision of these fast inhibitory-excitatory circuits.

Collectively, our analyses consistently implicate the high-gamma band in adult ADHD pathophysiology. Reduced network integration (longer CPL) in attention networks, diminished local processing efficiency (lower strength/clustering) in key DAN/VAN nodes, and widespread NBS-identified edge-level hypo-connectivity all converge within this high-frequency range. Gamma oscillations, broadly defined (approx. 30-100 Hz and higher), are increasingly recognized as vital for various cognitive processes, many of which are characteristically impaired in ADHD (Herrmann et al., 2004; Jensen et al., 2007). They are thought to reflect synchronous neuronal firing that enables efficient neuronal communication, feature binding, and the formation of transient neuronal assemblies necessary for active cognitive processing (Jensen et al., 2007). Specifically, gamma activity has been strongly linked to attentional selection, where it may mediate gain control for relevant stimuli and facilitate processing (Gruber et al., 1999; Jensen et al., 2007; Ray et al., 2008), with higher gamma activity associated with better attentional performance (Akimoto et al., 2014). Furthermore, gamma oscillations play a role in both working memory maintenance, potentially reflecting the active maintenance of information, and are predictive of successful long-term memory encoding and involved in retrieval (Herman et al., 2020; Herrmann et al., 2004). The fast temporal scale of gamma oscillations (10-30 ms periods) is proposed to allow for tighter temporal integration of neuronal inputs compared to slower rhythms, facilitating rapid information processing and long-range communication (Jensen et al., 2007). Given that deficits in attention, working memory, and often processing speed/efficiency are core features of ADHD, the consistent pattern of high-gamma disruptions observed in our study (reduced integration within attention networks, reduced local processing efficiency, widespread hypo-connectivity) points towards a potential fundamental impairment in the high-frequency network dynamics that support these very functions. The robust high-gamma findings underscore the value of MEG’s temporal resolution, revealing network dynamics potentially obscured in studies relying solely on the slower hemodynamic signals measured by fMRI. This specific high-frequency disruption may represent a core neurophysiological signature of adult ADHD.

### 4.2 Contrasting Alpha-Band Dynamics and Other Frequency-Specific Observations

Adults with ADHD showed increased integration (shorter CPL) in alpha band within the DMN compared to controls. This finding is notable given consistent reports of DMN dysfunction in ADHD (Castellanos et al., 2008; Fair et al., 2010; Franzen et al., 2013; Hoekzema et al., 2014; Uddin et al., 2008; Wilson et al., 2013) and might reflect an abnormal stabilization of an internally-focused state (Sonuga-Barke & Castellanos, 2007) or dysregulated DMN interactions (Hoekzema et al., 2014), observable with MEG’s direct neural measures.

Analysis of between-network communication, using NBS on average shortest paths, further revealed significantly shorter paths in the ADHD group within an alpha-band sub-network centered on the DAN. The pronounced difference between DAN and VAN suggests potentially less segregated or overly rigid communication between these attention systems in the alpha frequency range in ADHD. Typically, the DAN (mediating top-down control) and VAN (mediating bottom-up reorienting) exhibit relative segregation at rest, reflecting their distinct roles, but interact dynamically during tasks via specific anatomical and functional connector hubs (Spadone et al., 2015; Suo et al., 2021). Efficient between-network functional connectivity, particularly involving these hubs, has been linked to better cognitive control performance, such as motor inhibition (Hsu et al., 2020). Our finding of shorter alpha-band paths between DAN and VAN nodes in ADHD at rest could signify reduced segregation compared to controls. This might reflect a less differentiated or perhaps less mature network state (Farrant & Uddin, 2015), potentially hindering the flexible, context-dependent engagement and disengagement of these networks. While efficient communication between attention networks is crucial (Hsu et al., 2020), the specific increase in alpha-band coupling observed here might represent a form of maladaptive or overly rigid interaction, potentially interfering with, rather than facilitating, optimal attentional processing. This highlights how frequency-specific alterations in connectivity may yield different functional interpretations compared to broadband measures often used in fMRI.

Local graph analysis revealed further frequency-specific modifications. In the alpha band, the ADHD group showed increased node strength and clustering in the middle-anterior part of the left cingulate (part of VAN, anatomically overlapping with the cognitive division of the anterior cingulate cortex), a key region for executive functions often impaired in ADHD (Botvinick et al., 2004; Bush et al., 1999; Carter et al., 1998; Devinsky et al., 1995, 1995; Swick & Turken, 2002); this might reflect aberrant resting-state inhibition. Additionally, increased beta-band node strength in right somatomotor regions suggests altered baseline motor network processing (Stolk et al., 2019; Bu et al., 2020).

### 4.3 Contextualizing Findings: Relation to Prior Research

Our findings of widespread high-gamma hypo-connectivity in adults with ADHD contribute to a nascent MEG/EEG literature that has faced challenges of methodological heterogeneity and a predominant focus on pediatric populations (e.g. Khadmaoui et al., 2016; Janssen et al., 2017). Unlike some pediatric MEG studies reporting gamma hyperconnectivity, potentially due to developmental factors (Janssen et al., 2017), our source-space analysis using individual head models robustly identifies hypo-connectivity in adults. This resting-state observation may differ from task-related findings (Heinrichs-Graham et al., 2014) and, while a MEG study following individuals into young adulthood found mixed patterns including decreased DMN connectivity and increased connectivity between the DMN and other networks associated with persistent inattention (Sudre et al., 2017), our NBS results more broadly emphasize widespread reduced coupling within high-gamma networks involving DMN, DAN, and VAN nodes at rest. These findings align with reports of reduced resting-state gamma power in adult ADHD (Tombor et al., 2021) and broadly supports neurobiological models of large-scale network dysfunction (Castellanos & Proal, 2012; Kucyi et al., 2015; Sonuga-Barke & Castellanos, 2007), refining them by highlighting the critical role of disrupted high-frequency dynamics in the adult condition.

### 4.4 Predictive Insights, Methodological Considerations, and Future Outlook

Predictive modeling analyses indicated moderate potential for MEG-derived network metrics, especially high-gamma CPL (AUC ≈ 0.70–0.73), in differentiating adults with ADHD from controls. While not yet sufficient for standalone clinical diagnosis, the superior performance of high-gamma features underscores their significance and potential utility in future multimodal approaches, pending further validation. PLS-DA analysis further highlighted the weight of frontoparietal and ventral and dorsal attention networks connectivity in this high-gamma based group discrimination.

This study possesses several strengths, including MEG’s high temporal resolution for probing fast oscillations in the understudied adult ADHD population, a comprehensive graph-theoretical analysis, wPLI for robust connectivity, and use of a public dataset promoting reproducibility. Limitations include the cross-sectional design, modest sample size for heterogeneous ADHD, uncontrolled confounds in public data (e.g., medication), chosen parcellation/connectivity measure, and exclusive focus on intrinsic brain activity. Future research should prioritize replication in larger, clinically well-characterized adult cohorts, longitudinal designs, multimodal imaging (fMRI/DTI), and investigation of task-based connectivity and intervention effects.

## 5. Conclusion

In conclusion, this MEG study reveals significant, frequency-dependent alterations in functional brain network organization in adults with ADHD. The most consistent finding was widespread hypo-connectivity in the high-gamma frequency band, affecting integration within attention networks, local processing in key nodes, and specific connections primarily involving the dorsal attention, ventral attention, and default mode networks, with the left fusiform gyrus acting as a central hub in this hypo-connected system. Our findings advance the understanding of the neurophysiological basis of adult ADHD, highlighting specific alterations in high-frequency network communication that may be targeted in future diagnostic and therapeutic approaches.

## DATA AND CODE AVAILABILITY

The data used in this study is from the Open MEG Archive (OMEGA) accessible from https://www.mcgill.ca/bic/neuroinformatics/omega. The connectivity data and the code to reproduce all the findings of this study will be available upon request.

## AUTHOR CONTRIBUTIONS

PM: Conceived the study, performed data curation and analysis, prepared all visualizations, and wrote the initial manuscript. BTD: Provided critical revision and feedback on methods and manuscript, approved final version. MM: Provided critical revision and feedback on methods and manuscript, approved final version.

## ACKNOWLEDGEMENTS

M Moayedi is supported by a University of Toronto Centre for the Study of Pain — Pain Scientist Award, a Canada Research Chair (Tier 2) in Pain Neuroimaging, and the Bertha Rosenstadt Endowment Fund at the Faculty of Dentistry, University of Toronto. BT Dunkley is CSO at MYndspan Ltd and has received compensation for his industry work. The authors have no other conflicts to report.

