## Supplemental materials for "Functional connectivity differences in adult’s ADHD – a MEG study"

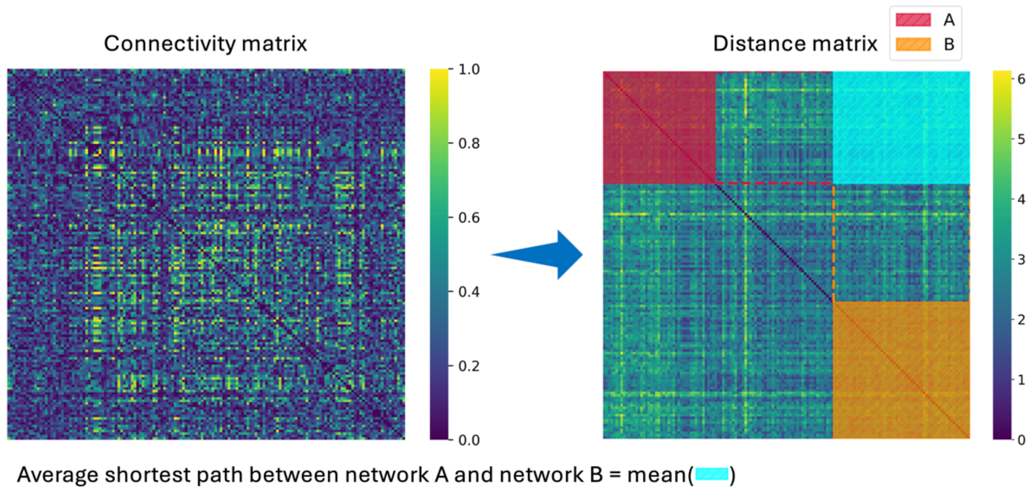


**Figure S1: Calculation of between network average shortest paths**. The distance matrix is calculated from the connectivity matrix using the Dijkstra's algorithm (Dijkstra, 1959), identifying the shortest path between each pair of regions. The between network average shortest path is then calculated by averaging the shortest paths between all the nodes of two networks of interest.

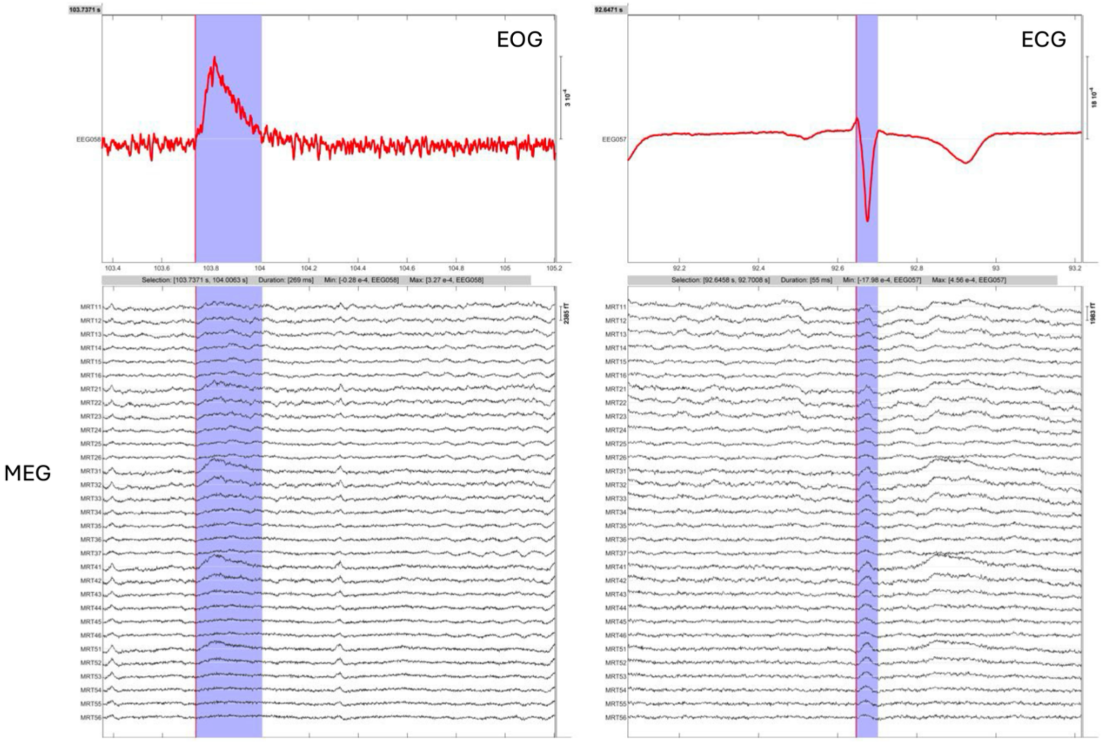


**Figure S2: The effect of eye blink (detected from the electrooculogram (EOG) signal) and heartbeat (detected from the electrocardiogram (ECG) signal) on the MEG signals.**

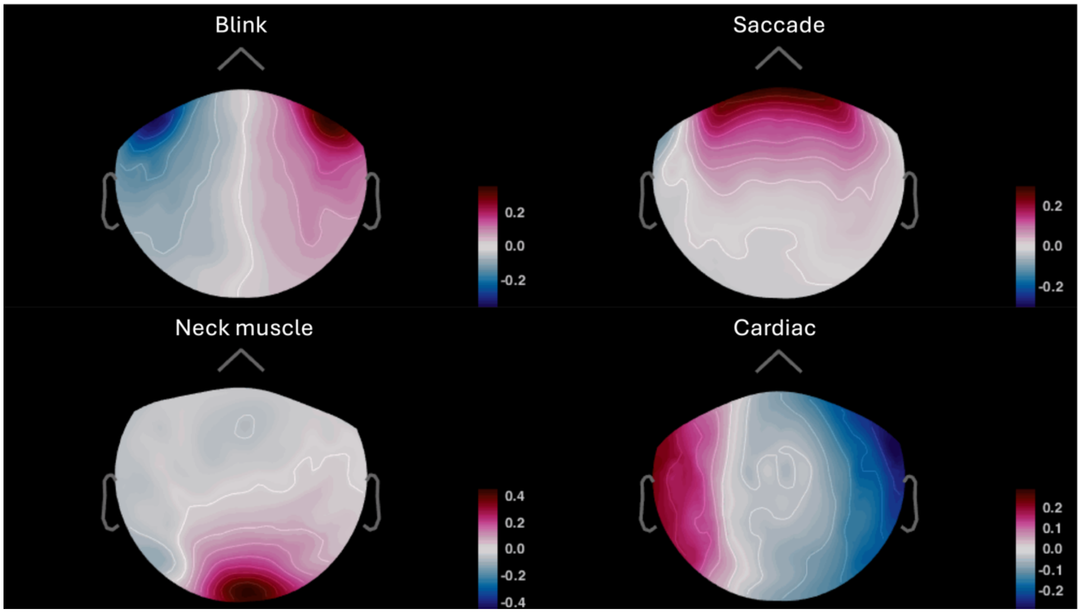


**Figure S3: Examples of artifacts components' topography.** The heartbeat and blink events were automatically detected from the ECG and EOG signals, respectively. The ECG and EOG signals were then inspected manually to confirm the events. Saccades were also detected manually by inspecting big shifts in the horizontal EOG and MEG signals. Periods of signal affected by neck muscle artifact are characterized by high frequency noise in posterior regions and were detected by visual. Saccade and neck muscle artifact periods defined manually but heartbeat and blink periods were defined by time windows of ±40 ms and ±200 ms around the detected events, respectively. SSP method was used to calculate principal components and remove those associated with artifacts. All the components were reviewed manually and artifactual components selected based on their explained variance and topography.
